# Shear Strain Stiffening in Cartilage is Governed by a Mechanical Phase Transition

**DOI:** 10.64898/2026.09.28.751808

**Authors:** Japheth Omonira, Joshua D. Clugston, Anna R. Barth, Stephen J. Thornton, James P. Sethna, Moumita Das, Lawrence J. Bonassar, Itai Cohen

**Author notes:** I.C. and L.B. designed research; Experiments performed and analyzed by J.O. and J.D.C. Network simulations done and analyzed by A.R.B. Written by J.O., J.D.C., A.R.B., L.B. and I.C. with all authors contributing. The authors declare that they have no competing interests.

## Abstract

Large shear deformation in cartilage can lead to tissue failure and injury, which are often precursors to diseases such as post-traumatic osteoarthritis. Under shear, cartilage strain-stiffens prior to failure, which makes stiffening an important indicator of changes to the loading of the extracellular matrix that gives rise to this tissue’s properties. While a variety of stiffening mechanisms ranging from uncrimping to fiber-alignment have been explored, whether such mechanisms apply to the depth dependent shear strain stiffening in articular cartilage remains poorly understood. Here, we present experiments and theory that support the idea that the strain-stiffening behavior is caused by a mechanical phase transition from a random isotropic network whose mechanics arise primarily from bending-dominated modes to one where the mechanics arise from stretching-dominated modes. Indeed, consistent with network model predictions, we find that stress-strain data for all depths can be collapsed onto a single universal curve that diverges at a depth-dependent critical strain. These results show that increasing the local concentration of collagen, which constitutes the primary load-bearing network in cartilage, decreases the critical strain at which the strain-stiffening transition occurs, while increasing the low-strain elastic modulus. Furthermore, we demonstrate that bulk compression leads to a higher critical strain and decreased low-strain elastic modulus. Network simulations capture this rich behavior, and indicate that under compression the network collapses, which allows for larger strains before the transition from bending-dominated to stretching-dominated mechanics occurs. Collectively, these findings shed light on the roles that tissue composition and axial strain play in the mechanisms that organize how articular cartilage stiffens during the lead-up to injury.

**Significance Statement:** We show that strain stiffening in cartilage, which is a precursor to tissue damage, is governed by a mechanical phase transition from a network whose properties arise from bending-dominated modes to one where the mechanics arise from stretching-dominated modes. This framework provides insights into the possible failure mechanisms that arise under extreme loading and points to new approaches for prevention and repair.

---

**A**rticular cartilage sustains decades of repetitive mechanical loading while maintaining exceptionally low friction and wear(1, 2). Despite this remarkable durability, cartilage is susceptible to acute mechanical insults(3, 4). Traumatic joint loading events, such as those associated with sports injuries, falls, or vehicle collisions, can impose extreme strains that exceed the tissue’s normal physiological range and initiate post-traumatic osteoarthritis (PTOA)(3–7). PTOA frequently develops even in the absence of overt tissue rupture at the time of injury, highlighting the importance of understanding the mechanical response of cartilage prior to catastrophic failure.

A defining feature of cartilage under large deformation is strain stiffening, in which the tissue becomes increasingly resistant to deformation as strain increases(8–10). This nonlinear behavior is widely observed in cartilage across loading modes(8, 11–14), and is often interpreted as a protective mechanism that limits excessive deformation. At the same time, strain stiffening precedes the onset of irreversible damage, suggesting that it marks a critical transition in how load is borne by the tissue. Understanding the physical origin of this transition is therefore essential for identifying how extreme loading drives cartilage toward disease and ultimately failure.

The mechanical behavior of cartilage arises from its extracellular matrix, which is composed primarily of two structurally distinct but mechanically interdependent networks(15). A network of collagen fibrils provides tensile strength and structural integrity, while a highly charged aggrecan matrix, anchored to hyaluronic acid, generates osmotic swelling stresses and resists volumetric deformation(15, 16). These constituents are distributed heterogeneously through the tissue depth, producing gradients in composition, microstructure, and linear mechanical properties. Near the articular surface, the collagen network is relatively sparse and highly oriented, whereas deeper regions contain denser and more isotropic collagen architectures accompanied by higher aggrecan concentrations(17). As a result, the local mechanical environment varies substantially with depth. Indeed, measurements of the linear shear modulus show a difference of one to two orders of magnitude between the compliant surface region and the more rigid deep zones of the tissue(18–21). Additionally, this heterogeneity in shear mechanics is further complicated by compressive loading. Under higher levels of compression, the surface region is even more compliant in shear, while the stiffness of deeper regions remains relatively constant(18). This difference in composition and mechanics as a function of depth raises the possibility that nonlinear responses, such as strain stiffening, could also be depth-dependent.

The mechanistic origins of cartilage’s shear strain-stiffening response have been a topic of interest for decades. Multiple models have been proposed to explain the strain-stiffening response in articular cartilage. A common phenomenological explanation of strain stiffening in collagenous tissues is the straightening of collagen crimps(22). Collagen fibrils exhibit a periodic, wave-like structure that is referred to as a crimp(23–25). When a load is applied, the initial soft “toe” region is caused by the straightening of these crimps without any stretching of the fibril. Once the fibril is completely straight, it begins to stretch, eliciting a stiffer “linear” region(26). In a fiber with fibrils varying in slack and crimp structure, individual fibrils will fully straighten at different amounts of stretch, causing a progressive recruitment of crimped fibrils to straightened fibrils that collectively results in the “heel” region as the fiber transitions from soft to stiff(27). The collagen uncrimping and fiber recruitment process has been used to describe the nonlinear mechanical behavior of collagenous tissues such as tendon, skin, and eye(28–32). The parameters these crimping models use often include material properties, such as crimp angle and fiber diameter, and mechanical properties such as the stiffness of the crimped fiber, the stiffness of the uncrimped fiber, and a characteristic (or critical) strain (or stretch) signifying the strain at which the fiber becomes fully uncrimped. While the relationship between fiber diameter and the modulus in both the crimped and uncrimped state has been investigated(26, 33–35), whether the threshold for uncrimping or strain stiffening depends on concentration or volume fraction of collagen remains poorly understood. Analysis of the models for the transition between the crimped regime and uncrimped regime for collagen suggests that to first order, the critical strain where this transition occurs should be independent of the collagen concentration or fiber size. Instead, these models suggest that with increased fiber size, the range of strains over which the transition occurs should become broader. Therefore, measuring whether the mean critical strain associated with stiffening depends on concentration and determining whether the strain distribution broadens for thicker fibers associated with the deep zone would enable us to determine the applicability of this model as a mechanism for shear strain stiffening in articular cartilage.

Strain-stiffening in soft biological tissues has also been explained using the worm-like chain (WLC) model, which is often used in the study of semiflexible polymers stiff enough to retain a straight conformation but flexible enough to allow thermal fluctuations, such as DNA and F-actin(36–38). In this model, a polymer is treated as either a continuous string or a discretized chain of segments whose unstrained shape is determined by a persistence length that depends on temperature and the polymer’s bending stiffness(39, 40). If the persistence length is much larger than the polymer length (high bending stiffness and low temperature), the polymer is very stiff and too inflexible to bend. If the persistence length is much smaller (high temperature and low bending stiffness), the polymer is very soft and too flexible to maintain a straight conformation. When the persistence length is on the order of the polymer length, the polymer is semiflexible, and entropic contributions to the energy must be considered(41). As a bent polymer straightens under tension, the polymer transitions from random to oriented states, and its stiffness increases dramatically due to the high stretching stiffness of the polymer(39). In other words, strain stiffening in the WLC model is an entropic process caused by the straightening out of thermal fluctuations in the polymer. Importantly, the collagen fibers making up the extracellular network in articular cartilage are athermal(38, 42, 43). Thus, for this model to be applicable, the interplay between the collagen fibers and the surrounding aggrecan molecules would need to provide an effective temperature that leads to fluctuations in the collagen fiber conformations. In this case, the critical stress associated with the stiffening transition should be dependent on the effective temperature, where increasing the effective temperature results in an increase in the critical stress. Importantly, the WLC model is a single fiber model and, as such, predicts a single mean critical strain independent of collagen concentration. Analysis of multiple fibers with a distribution of initial states would broaden the range over which the transition occurs, but not affect the mean critical strain. As such, determining whether the critical strain depends on collagen concentration would be useful for determining the applicability of this model for strain stiffening in cartilage.

Beyond individual fiber models, it has been proposed that the realignment of collagen fibers under increasing shear strain can lead to stiffening(44). These models hold that when strained, fibers are under a tensile load that first causes them to align in the direction of loading and then stretch(44, 45). For bulk materials, the tissue stiffness depends on the initial distribution of fiber orientation. Tissue with fibers already aligned in the direction of the load should require less strain to fully align and stiffen the tissue, while isotropic material would require more strain to align the fibers and stiffen(45). Cartilage tissue is primarily comprised of regions with a heterogeneous fiber alignment throughout the depth of the tissue. The articular surface has a small ~ 50 µm region where the fibers are aligned parallel to the surface. Deeper regions have a more isotropic orientation(17). Accordingly, fiber alignment models predict that the highly aligned superficial zone should have a small critical strain while the isotropic middle zone should have a larger, relatively unchanging critical strain. Importantly, such models predict a critical strain that depends on fiber orientation but is independent of fiber concentration.

Some have proposed combining the fiber alignment model with the collagen crimping model. In that case, fiber realignment would be followed by elongation of the fibers due to uncrimping and then fiber stretching. This suggests that an individual fiber has three strain-related moduli regimes, one associated with the force needed to align the fiber, one associated with uncrimping the fiber, and one associated with stretching the fiber(44, 46). The addition of uncrimping to this model should broaden the range of critical strains over which the transition occurs, but preserves the dependence of the critical strain on fiber orientation and the lack of dependence on collagen concentration. As such, the applicability of fiber alignment models on strain-stiffening in articular cartilage can be verified by measuring whether the critical strain is dependent on concentration or collagen fiber alignment.

To take into account the architecture of collagen based extracellular matrices, it may also be useful to investigate the applicability of network models to strain stiffening in cartilage. Such models consist of 2-D or 3-D disordered athermal networks of nodes and bonds, which are characterized by bending and stretching moduli. When the network transitions from a bond bending dominated regime to a bond stretching dominated regime, it exhibits characteristics typically associated with thermodynamic phase transitions. For example, near the critical strain associated with the bending to stretching transition, the stress obeys power law scaling with a universal exponent(47). This prediction was subsequently rederived using field theory (48) and numerically verified (49). Interestingly, the dependence of this mechanical phase transition on the number of fibers, or concentration, is still poorly understood. Because adding more fibers drives unstrained networks closer to stretching rigidity, it is expected that increasing the concentration of bonds would result in smaller critical strains. Therefore, network models are expected to show a dependence of the critical strain on bond occupation probability. If such models apply to strain stiffening in cartilage, we would expect to observe a dependence of the strain associated with stiffening on the collagen concentration.

Finally, we note that there have been several attempts to fully capture the behavior of cartilage under both compression and shear. For example, constitutive models of articular cartilage mechanics have evolved from a linear elastic model in 1972(50), to a biphasic model in 1980(11), to now fibril-reinforced multiphase models(51–54). In fibril-reinforced models, a fibrillar phase representing collagen fibrils is used to capture the nonlinear mechanical behavior of cartilage under shear(52, 55, 56). Thus, while such models introduce additional mechanisms that capture, for example, the poroelastic behavior of cartilage under compression, the shear strain stiffening behaviors in these models arise from the previously described mechanisms.

Remarkably, while the bulk or average shear strain-stiffening response in cartilage has been the subject of experimental investigations for decades, its dependence on structural and mechanical features that vary with depth remains poorly understood. Results from tensile strain measurements of sectioned tissue indicate that tensile stiffening depends on zonal architecture(8, 57). Additionally, prior work has established that homogeneous collagen networks, aggrecan gels, and composite fibrous materials can each exhibit nonlinear behaviors under shear when these systems shift from bending-dominated to stretching-dominated mechanics (58–67). However, how such mechanisms interact to produce the depth-dependent stiffening response in tissues such as cartilage, particularly under large shear stresses relevant to traumatic loading, remains poorly understood. While all the mechanisms described above propose that stiffening arises when the fibers move from a bending dominated to a stretching dominated regime, it is unclear which mechanisms are most relevant. Importantly, the various mechanisms have different predictions for how the critical strain associated with the transition depends on the collagen concentration. Therefore, to distinguish between these mechanisms, it is critical to measure how the shear strain stiffening transition depends on depth-dependent composition and proximity to mechanical thresholds within the extracellular matrix.

Here, we use state-of-the-art confocal elastography and FTIR microscopy measurements in combination with fiber network simulations to investigate the dependence of shear strain stiffening in articular cartilage on depth and composition. We find that the threshold strain for the stiffening transition does indeed vary with depth and depends on collagen concentration. Further, we find that the stiffening data for all depths collapse onto a universal curve that diverges at a critical strain that varies with depth. This collapse provides compelling evidence that stiffening in cartilage can also be thought of as a mechanical phase transition. Additionally, we found that the critical strain and the shear modulus in the linear regime prior to stiffening depend on tissue composition and compression. Surface regions with lower collagen concentrations had lower linear shear moduli and higher critical strain values, while deeper regions with higher collagen concentrations had higher linear shear moduli and lower critical strains. These results are consistent with data collected from fiber network simulations that used varying bond occupation probability as a surrogate for collagen concentration. Finally, we show in both experiments and simulations that increasing the level of applied compression prior to shear resulted in lower linear shear moduli and higher critical strains. Collectively, these results strongly suggest that shear strain stiffening in cartilage is best described by the mechanical phase transition mechanism.

By linking nonlinear mechanical response to underlying network architecture, our results provide a physical framework for understanding how cartilage stiffens, protects itself, and ultimately becomes vulnerable to damage under extreme loading. As such, this framework offers new insight into the mechanical precursors of PTOA and suggests design principles for strategies aimed at preventing or repairing cartilage damage.

## Materials and Methods

### Sample Preparation

One-to-three-day-old neonatal bovine knee joints were acquired from a local abattoir (Gold Medal Packing, Rome, NY). Neonatal cartilage was chosen because its shear modulus profile is similar to that of human cartilage and because it is easy to obtain, readily available, and used in numerous prior studies(19, 68). Cylindrical explants 6-mm in diameter were extracted from the medial and lateral femoral condyles, trimmed to 3 mm thickness, and split in half to create two hemicylindrical samples. One half was stored in phosphate-buffered saline (PBS) until mechanical testing; the other half was fixed in 10% PBS-buffered formalin for compositional and structural measurements.

### Mechanical Testing

To prepare for mechanical testing, each sample was stained for 1 h in PBS with 7 mg/mL 5-dichlorotriazinylaminofluorescein (5-DTAF) (Life Technologies, Carlsbad, CA). The samples were then rinsed with PBS for at least 30 minutes to remove any excess dye. The dyed sample was loaded into a Tissue Deformation Imaging Stage (TDIS) (Harrick Scientific, Pleasantville, NY). The TDIS consists of two plates, a shearing plate and a nearly stationary force measurement plate, whose relative positions can be adjusted using three micrometer screws. A piezo motor, enabling dynamic unidirectional motion of the shearing plate was used to control the applied strain. The sample was affixed to the shearing plate and the stationary plate using cyanoacrylate glue to ensure no slip at either end of the tissue, with the articular surface attached to the moving shear plate. While mounted into the TDIS, the sample was kept in a PBS bath to maintain tissue hydration during testing(19, 20).

The TDIS apparatus was then mounted onto an inverted spinning disk confocal microscope (3i Marianas). The tissue was first placed under a static compressive load, and allowed to reach osmotic equilibrium. Then, an oscillatory shear strain ranging from ~ 1% to ~ 70% peak amplitude and either 1 Hz or 0.01 Hz frequency was applied using the piezo motor controlled by LabView software(19).

The tissue was imaged using a 10x objective at approximately 20 frames per second. The resultant videos were analyzed using automated computer tracking of the displacement throughout the depth of the ~3 mm sample. The displacement of the tissue was automatically tracked using the MatPIV(69) MATLAB script. The local shear strain, *γ*, with a depthwise resolution of 10.4 µm was calculated from the displacement by differentiating the displacement with respect to the depth. The force applied to the tissue was calculated by observing the displacement of the nearly stationary shearing plate whose stiffness was calibrated prior to the experiments. The local shear stress, *σ*, was calculated by dividing this force by the cross-sectional area of the tissue, which is assumed constant throughout the depth of the tissue (20).

### Fourier-Transform Infrared Microscopy

Relative collagen and aggrecan concentrations were measured using FTIR-I in the same manner described in previous studies(21, 70–72). The second half of the bisected sample was sectioned into 4-*µ*m thick samples, which were placed on 2-mm-thick *BaF*_2_ slides. The sections were dewaxed in three xylene baths for two minutes each, and rehydrated in three baths of ethyl alcohol (100, 95, and 70%) for two minutes each. Samples were loaded into a Hyperion 2000 Fourier Transform Infra-Red Imaging (FTIR-I) microscope (Bruker, Billerica, MA.) in transmission mode and data were collected for wave numbers between 600 and 4000 *cm*^*−*1^ at a resolution of 4 *cm*^*−*1^. A 15x objective was used to acquire spectra throughout the depth of the tissue, with a slit aperture configured to acquire spectra over a rectangular region 25 µm x 200 µm, where the long dimension was parallel to the articular surface. Fifteen background corrected scans along the tissue depth at 25 µm intervals were acquired for each sample and averaged to generate each IR spectrum. This procedure was repeated three times for each tissue sample with each scan separated laterally by roughly 2 mm taking care to avoid blood vessels and other artifacts.

Using Beer-Lambert’s law, the relative concentrations of aggrecan and collagen were calculated from the acquired spectra. Infrared absorbance is proportional to molecular concentration. By fitting a linear combination of previously measured spectra for aggrecan and collagen to the measured spectra of the samples, assuming the contributions of other extracellular constituents to be negligible, we determined the relative concentrations of the constituents(70).

The depth-dependent concentrations found for each sample were then paired with the other half of the bisected explant. To ensure that we are correlating the FTIR-I measurements on uncompressed samples with the appropriate region in the compressed and sheared sample, we keep track of the amount of tissue compression during axial loading and refer to each depth according to the uncompressed concentration. This procedure also allows us to take into account the fact that compression does not alter the network connectivity responsible for stiffening. To track the depth-dependent compression, we matched the location of prominent features of the tissue under an axial load to those features’ locations in the uncompressed sample. By tracking such features throughout the depth of the tissue, we obtained values for the compression with a resolution of *~*50-100 µm with higher resolution near the articular surface and lower resolution deeper in the tissue. Finally, FTIR-I data were paired with the appropriate tissue depth in the mechanically tested samples.

### Biochemical Assay

Biochemical assays of the mechanically tested half of the explant were performed to calibrate the FTIR data and determine the absolute concentrations of collagen and aggrecan. Briefly, light absorption measurements read on a microplate reader (Biotek Synergy HT) of digested samples were compared to a serial dilution standard curve made for each assay. The digested samples were tested in triplicate for statistical analysis.

To digest the samples, they were first weighed to obtain a wet weight, then frozen at −80^*°*^ C before being dehydrated in a lyophilizer for 48 hours and weighed again to obtain their dry weight. The samples were then incubated in papain digest buffer (PDB) containing 0.125 mg/mL papain (Sigma) and 10 mM N-acetylcysteine (Sigma) in PBE buffer (100 mM phosphate and 10 mM EDTA, pH 6.5) for 12 hours at 60^*°*^C.

Serial dilution standards were created to make a standard curve from which the unknown concentration of the samples can be deduced. Chondroitin sulfate (C-6-S) was used as the standard in the glycosaminoglycan (GAG) biochemical assay to determine aggrecan concentrations. Twenty-five milligrams of C-6-S were dissolved in one milliliter of PDB. 100 µL of the 25 mg/mL solution was added to 9.9 mL of PDB, to make a 250 µg/mL solution. This solution was used to make seven standards via 1:2 serial dilutions, resulting in standards that ranged from 125 µg/mL to 1.95 µg/mL. A 0 µg/mL standard was included.

The absorption measurements were done on a 96-well plate. Fifty microliters of the digested samples were added to individual wells in triplicate. Fifty microliters of the C-6-S standards were also loaded into individual wells in triplicate. Five microliters of alginate lyase buffer was added to each well, followed by two hundred fifty microliters of DMMB dye (Sigma-Aldrich, pH 3). The plate was shaken for 30 seconds before absorbance was read using the microplate reader with an excitation wavelength of 525 nm.

Hydroxyproline was used as the standard for the biochemical assay to determine collagen concentrations. Collagen is approximately 13.5% hydroxyproline, so the end measurement is multiplied by 7.4. The standards were made by dissolving 10 mg hydroxyproline in 1 ml PDB to make a 10 mg/mL solution. This solution was used to make nine standards via 1:2 serial dilutions ranging from 10 mg/mL to 19.5 µg/mL. A 0 µg/mL standard was included.

Digested samples and hydroxyproline standards were further prepared for the biochemical assay in triplicate. A 50 µL aliquot of each sample or standard was added to an individual tube, followed by 50 µL of 2 M NaOH. The tubes were heated at 110^*°*^C for 18 hours. The following solutions were then added sequentially, with vortexing after each addition: 30.5 µL of HCl solution, 100 µL of 0.01 M CuSO_4_, 100 µL of 2.5 M NaOH, and 100 µL of 6% H_2_O_2_. The samples and standards were incubated at room temperature for 2 hours, vortexed, heated at 80^*°*^C for 5 minutes, and then frozen for 4 minutes. Next, 400 µL of 1.5 M H_2_SO_4_ was added to each tube, followed by a second 5-minute freezing step at −80^*°*^C. A 200 µL volume of DMAB solution (0.5 g DMAB in 10 mL propanol) was then added to each tube. The tubes were vortexed and heated at 70^*°*^C for 15 minutes. Finally, 200 µL aliquots of each sample and standard were transferred to a 96-well plate, and absorbance was measured at 540 nm.

To correlate the concentrations of collagen and aggrecan from the biochemical assay to the FTIR compositional measurements, the average concentrations of the constituents were determined by dividing the total concentrations ascertained from the biochemical assays by the volume of the sample, calculated by multiplying the thickness of the sample by its cross-sectional area. The average concentration was then used to calibrate the relative concentrations found using FTIR-I. Pearson correlation coefficients were calculated within each sample–compression dataset across matched tissue depths and are reported as the mean ± SD across all datasets.

## Results

### Depth-dependent shear strain stiffening

Relating the compositional and structural characteristics to the shear stiffening behavior of cartilage tissue requires matched data sets for the mechanics and composition. In particular, articular cartilage exhibits a spatial heterogeneity in collagen concentration, aggrecan concentration, fiber orientation, and shear modulus(15, 17, 18, 73–75). Therefore, we collected mechanical and compositional data for 10 neonatal bovine condyles, combining FTIR imaging and confocal elastography on matched bisected samples. Confocal elastography was used to measure the local tissue mechanics while FTIR was used to measure local tissue composition.

To extract the depth-dependent tissue mechanical properties, we used confocal elastography to track the displacement of bovine cartilage under static compression and large amplitude quasistatic oscillatory shear (see Materials and Methods section). Example micrographs of stained cartilage tissue under 41% axial compression and various applied shear strains are shown in Fig. 1a. The photobleached lines are used to assess the level of shear strain. The top row shows three micrographs corresponding to bulk shear strains of 0%, 0.5%, and 1.0% relative to the uncompressed thickness, while the bottom row shows the same sample with bulk strains of 0%, 5%, and 10%. The images show that under low shear strain, the photobleached lines primarily bend in the surface region. This localized response, which has been reported previously (18, 21, 76), indicates that the strain in the tissue is concentrated in the surface region which is more compliant. For larger strains, the photobleached lines indicate that the strained region extends into the tissue bulk. By analyzing the slopes of the photobleached lines, we extract the local strain, *γ*, as a function of depth. The shear stress *σ* was determined by measuring the force corresponding to the deflection of the nearly static shearing plate and dividing by the surface cross section of the tissue. Importantly, the tissue samples were cut such that the cross-sectional area and therefore the stress was constant throughout the depth of the tissue. To obtain the relationship between stress and strain, samples were continuously subjected to the quasistatic oscillatory shear while sufficient data were collected at each depth to obtain a full cycle or Lissajous curve.

**Fig. 1.**
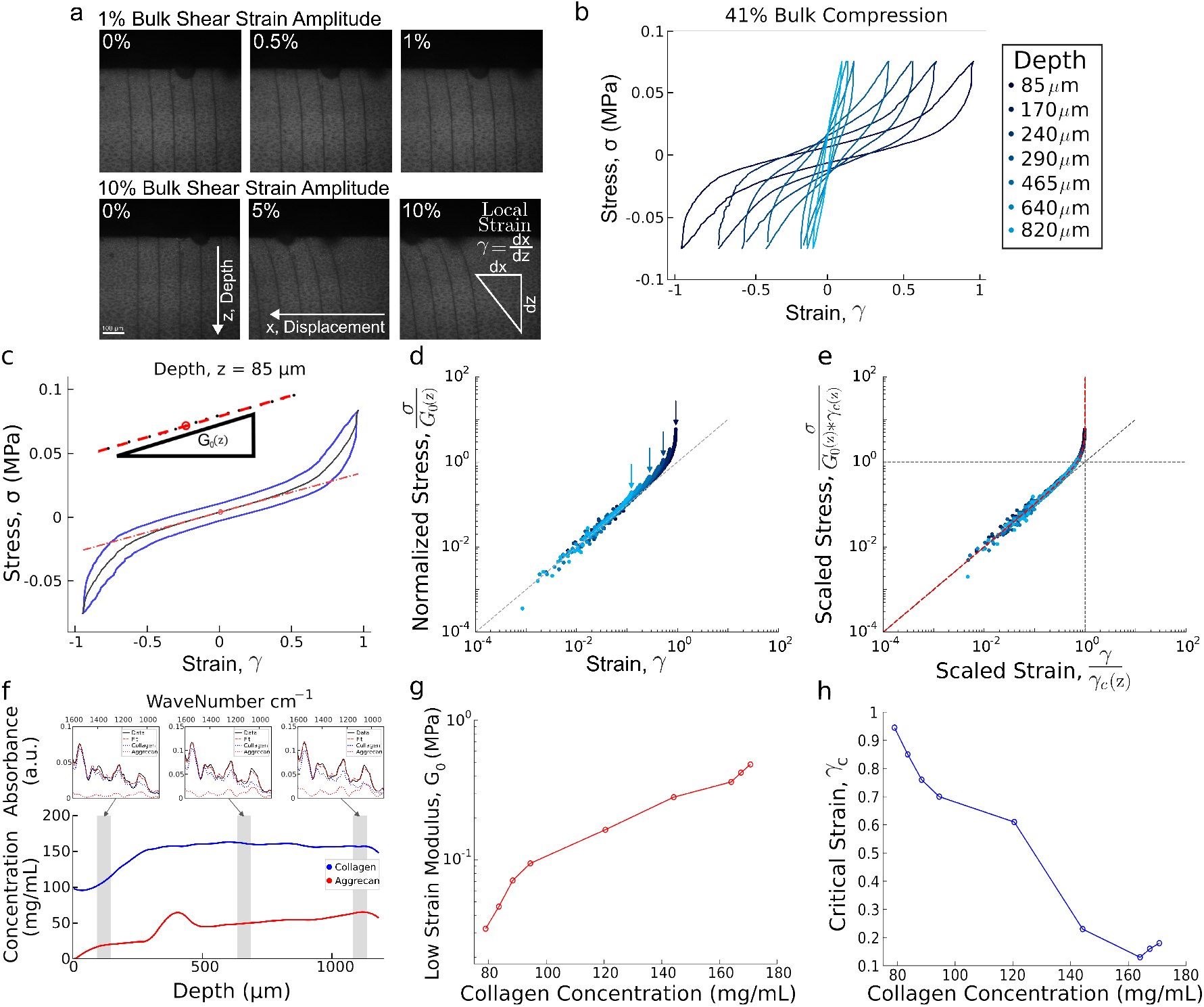
Analysis of the mechanical and compositional properties. a) Confocal micrographs taken of articular cartilage under varying levels of shear. b) Stress-strain Lissajous curves for 7 representative tissue depths at 41% bulk compression. Near the articular surface is colored dark blue and deeper tissue regions are colored light blue. c) The elastic stress is estimated for the Lissajous curves. The tangent slope at zero strain is *G*_0_, the zero-strain modulus. d) The elastic stress is normalized by *G*_0_. The normalized stress for each depth falls on a line of slope 1 at low strain and deviates from a linear response at differing values of strain. e) Scaling collapse of the stress-strain curves. Data that fall along the dashed line of slope 1 indicate a linear mechanical response. When the tissue stiffens at higher strain, the data deviate from the linear response. This deviation follows the Σ function (red dashed line). f) Collagen and aggrecan concentrations vs. depth for a representative sample. g) Values of *G*_0_ versus collagen concentration. h) Values of *γ*_*c*_ versus collagen concentration.

These data were used to determine the dependence of strain stiffening on depth. A typical sample’s Lissajous curves at seven representative depths are shown in Fig. 1b. We find that regions deeper in the tissue exhibit typical elliptical Lissajous curves indicating a nearly linear viscoelastic response. The high slopes, which correspond to the tangent modulus *G*_*T*_ = *dσ/dγ*, indicate these regions have a modulus approaching ~1 MPa. In contrast, the regions near the tissue surface have a lower slope at small *γ*, indicating that they are much more compliant with moduli approaching ~30 kPa, but rapidly increase in slope or stiffen by over an order of magnitude, at higher local strains, *γ*. We find that the local strain corresponding to the onset of stiffening behavior decreases with depth. Collectively, these data indicate that stiffening in cartilage from a compliant network at low strain to a stiff network at high strain is highly depth dependent.

Prior studies have shown that collagen fibrils in cartilage are only highly aligned in the ~50 µm region near the articular surface(71). The remaining tissue regions are characterized by a relatively isotropic distribution of fiber orientations. Such studies have shown that the broad range of tissue mechanics (in the linear regime) primarily arises from variation in the fiber concentration(71). Because the uncrimping, the fiber alignment, and the worm like chain models are expected to depend sensitively on fiber alignment, and/or are independent of fiber concentration, our observations rule out these mechanisms. As such, we focus the remaining investigation on the mechanical phase transition arising from network models for shear strain stiffening.

### Collapse of local shear stress on a single universal curve

A key prediction of network models is that the transition from the compliant to stiff regime arises from a mechanical phase transition, and that characteristic scalings should emerge at the onset of stiffening that are insensitive to the specific interactions of the network constituents. Inspired by these models, we expect that when properly scaled, strain stiffening data for the different depths should collapse onto a single universal curve. To test for this collapse, we first normalized the local stress by the zero-strain elastic modulus, *G*_0_. To determine *G*_0_, the elastic contribution to the stress was approximated by finding the midpoints between the upper and lower halves of the stress-strain curves. The value of *G*_0_ is then determined by the tangent of the midline at zero strain (Fig. 1c). We then divided the stress by the linear shear modulus *σ/G*_0_(*z*) to scale out the modulus in the linear regime (Fig. 1d) and find that the data collapse at low strains and deviate from the collapse at higher strains. The deviation from the linear behavior at large strains indicates shear stiffening. Furthermore, we find that the deviation from linearity is depth dependent. Next, we scaled both the normalized stress and the local strain *γ* by the critical strain, *γ*_*c*_, which is the strain at which the data would diverge, were it to follow a power law to its asymptote. The best fit to the data (see supplementary information for fitting procedure) indicates a power-law exponent of 0.53 ± 0.07, consistent with the mean-field prediction of 0.5 (47–49). We find that the data at all depths collapse onto a single curve (Fig. 1e), providing strong evidence that the shear strain-stiffening behavior of articular cartilage is governed by a mechanical phase transition.

### Correlation of strain-stiffening behavior to local composition

To relate the local mechanical data to composition, we determined the local collagen and aggrecan concentrations using FTIR-I and biochemical assays. FTIR-I enabled us to determine the relative amounts of aggrecan and collagen throughout the depth of the tissue while biochemical assays for GAG and hydroxyproline were used to calibrate the FTIR data to the average concentration of the sample. A representative sample is shown in Fig. 1f. The three absorbance spectra at the top show the measured spectra and the linear fit to that data at the three different positions in the tissue, the depths of which are shown in the bottom graph by the gray bars. The bottom plot shows the concentrations of aggrecan and collagen as a function of depth. Consistent with previous studies, we find lower concentrations of collagen and aggrecan near the articular surface and an increased concentration in deeper regions of the tissue. Since collagen is the primary load bearing network, we plot the low strain modulus G_0_ and critical strain *γ*_*c*_ versus collagen concentration in Figs. 1g,h (a similar analysis for the dependence on aggrecan is presented in the supplementary materials). We find that the mechanical data are highly correlated to the collagen concentration where the low strain modulus increases rapidly with collagen concentration (correlation coefficient 0.91 ± 0.11) while the critical strain decreases with collagen concentration (correlation coefficient −0.66 ± 0.30). As such, these data suggest a strong dependence of the mechanical phase transition on the underlying network composition.

### Strain stiffening in network simulations

To test if the network models can capture the observed scaling and dependence on collagen composition, we investigated these phase transitions in simulated networks with different bond occupation probabilities. Here, the bond occupation probability was used as a surrogate for collagen concentration. We note, however, that this is a gross simplification and that it is likely that other factors including the fiber size distribution, connectivity, and crosslinking may all contribute to this model parameter. For the simulations, we generated 2D triangular lattices with *N* = 1080 nodes and periodic boundary conditions in both directions. The lattices are occupied by two networks: a fully-occupied weak subnetwork with stretching modulus *µ*_*w*_, and a partially-occupied strong subnetwork with stretching modulus *µ*_*s*_ and bond occupation probability *p*. Here, the weak subnetwork provides elasticity in the low-strain limit and models elastic contributions from the bending of the collagen fibers, cooperative effects with aggrecan, and other extracellular matrix constituents. The strong subnetwork is meant to model the elastic contributions due to stretching of collagen fibers at high strain. The relative stiffness was *µ*_*w*_ */µ*_*s*_ = 10^*−*4^. We varied *p* from 0.4 to 0.6, which is a range where the strong subnetwork is above connectivity percolation but below rigidity percolation, and where significant strain stiffening is observed in the model.

To measure the shear modulus of these networks, we applied a simple shear strain *γ*. The lattices are deformed by applying the proper transformation matrix to both the nodes and the boundaries (77) (under a shear deformation, this procedure is equivalent to Lees-Edwards boundary conditions). At each strain step, the network’s energy is calculated according to

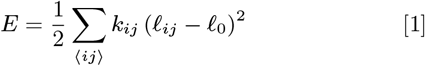

where the sum is over adjacent nodes indexed by *i* and *j, *l**_*ij*_ is the distance between nodes *i* and *j, *l**_0_ is the bond rest length, and *k*_*ij*_ is the stiffness of the bond between the nodes. Note that *k*_*ij*_ may be 0, *µ*_*w*_, or *µ*_*s*_ + *µ*_*w*_, depending on the occupation of the bond between nodes *i* and *j*. Unlike similar network simulations which often use an approximate form of this energy valid for small displacements, here we explicitly calculate the stretched length of each bond in order to capture the response of the network even at high strains, where node displacements might be large.

We probed the strain-dependent response of the networks by incrementally increasing *γ*. At each strain step, we relaxed the node positions to minimize Equation 1 using a trust region method and we tracked the resultant equilibrium energy *E*(*γ*) as the applied strain is increased. For each value of *p*, we averaged over 50 realizations of randomly occupied strong subnetworks to obtain the average energy ⟨*E*(*γ*) ⟩. The resulting differential stress, calculated as *σ* = *d* ⟨*E* ⟩ */dγ*, is shown in Fig. 2b. Consistent with the experimental data, the networks exhibit linear elastic behavior at low strains and strain-stiffen at higher strains.

**Fig. 2.**
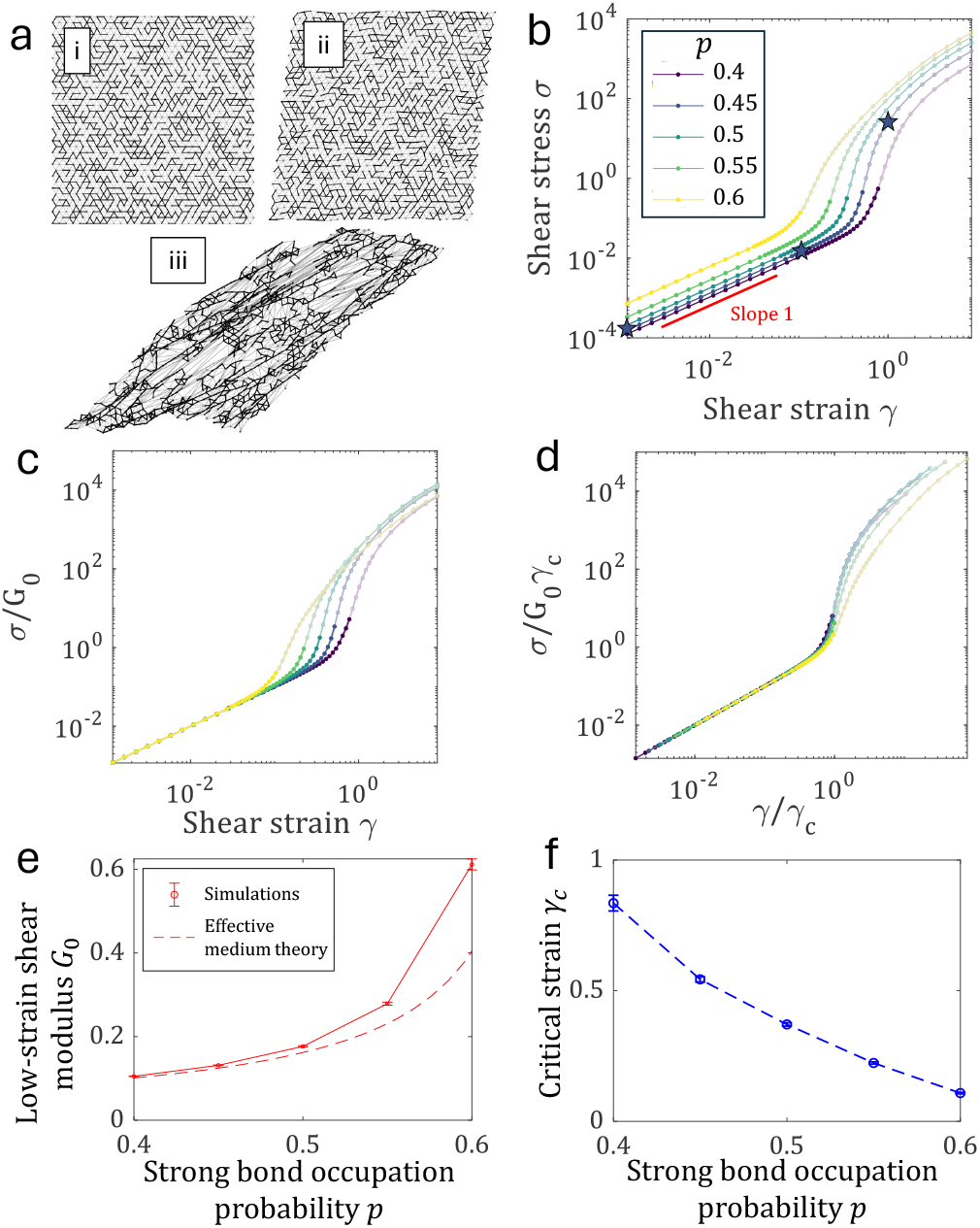
Simulations of double-networks with a fully-occupied weak subnetwork and a randomly-occupied strong subnetwork exhibit strain stiffening. (a) Images of a single simulation under increasing shear strain. (b) Stress *σ* versus strain *γ* in network simulations with different bond occupation probabilities for the strong subnetwork. The networks exhibit a strain-stiffening transition. The three stars correspond to subplots (a,i), (a,ii), and (a,iii) respectively. The highlighted data show points below the critical strain (*γ < γ*_*c*_). The red line shows a slope of 1, which corresponds to the networks’ linear elastic response at low strains. Error bars are shown as semi-transparent backgrounds behind each line of data. (c) *σ/G*_0_ plotted against strain *γ* shows a good collapse of the data at low strains, but each curve exhibits a different critical strain *γ*_*c*_. (d) Scaled stress *σ/*(*G*_0_*γ*_*c*_) plotted against scaled strain *γ/γ*_*c*_ shows a good collapse of the data for strains below the critical strain (*γ/γ*_*c*_ ≲ 1). (e) Low-strain elastic modulus increases as a function of strong subnetwork bond occupation probability. (f) Critical strain decreases as a function of strong subnetwork bond occupation probability.

To demonstrate that the simulation data exhibit universal behaviors similar to those measured in the experiments, we repeated the scaling analysis. We first examined the linear response of the simulated networks to determine the low-strain modulus *G*_0_. The linear response regime appears as a straight line of slope 1 on log-log axes, as shown in Fig. 2b. We observe that the low-strain modulus increases as a function of *p*, which can be seen as an upward shift of the linear regime in Fig. 2b. To quantify the dependence of the low-strain modulus on *p*, we simply differentiated the stress at the lowest strain in the simulations, 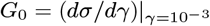. To scale out the low strain modulus, we divided the stress by *G*_0_ and plotted *σ/G*_0_ versus strain in Fig. 2c. As with the experimental data, the data collapses at low strains and deviates from the collapse at higher strains associated with stiffening. Further, we find that as the bond occupation probability increases, the onset of strain stiffening occurs at lower strains.

To further collapse the data, rather than empirically shifting the curves to obtain the best collapse and then fit the data, we determine the critical strain directly. Specifically, to determine the critical strain *γ*_*c*_(*p*) we ran another set of simulations with no weak subnetwork, as the critical strain does not depend on the weak subnetwork stiffness (47, 48). In the case where there is no weak subnetwork, the critical strain is simply the strain at which the network becomes rigid. For each value of *p* we randomly generated 50 networks with no weak subnetwork. For each network, we determined the critical strain *γ*_*c*_ by finding the strain at which *E* rises above the numerical precision of our simulations, indicating the network has become rigid. We then averaged these values to obtain *γ*_*c*_(*p*). Finally, we normalize the simulated networks’ stress and strain by this critical strain, plotting *σ/*(*G*_0_*γ*_*c*_) versus *γ/γ*_*c*_, and observe a collapse of the data up to strains of *γ/γ*_*c*_ ≈ 1, as shown in Fig. 2d. This collapse is consistent with the collapse of the data observed in the cartilage experiments. Furthermore, these network simulations exhibit power law scaling of the stress similar to that observed in the experimental data. Past studies of similar undercoordinated strain stiffening networks have shown that these models strain stiffen with a scaling *σ* ~ (*γ*_*c*_ *− γ*)^*−δ*^, where *δ* = 1*/*2 in the mean field limit (47, 48). Some variation in *δ* is observed in our simulations where the mean field theory does not apply; see the Supplemental Materials. The mean field value *δ* = 1*/*2 is well within the range measured in the experimental system.

Finally, we used these scaling analyses to extract the dependence of the low strain shear modulus *G*_0_ and the critical strain *γ*_*c*_ on the bond occupation probability. Consistent with the experiments, we observe that the low-strain modulus increases with the strong subnetwork bond occupation probability *p* (Fig. 2e). We find that this trend is captured by an effective medium theory (described in the Supplementary Materials). Additionally, we find that the critical strain decreases with the strong subnetwork bond occupation probability (Fig. 2f). Collectively, we find that both the qualitative dependencies on bond occupation probability and the quantitative prediction for a mean field exponent of ~ 0.5 describing the diverging stress are consistent with the experimental data.

### Strain stiffening under compression

Under physiological conditions, cartilage is often under axial compression, which should affect the depth-dependent stiffening behavior. Specifically, compression has been shown to make cartilage more compliant presumably due to fiber buckling, and this buckling should extend the regime of linear response so that stiffening occurs at higher critical strains. To conduct our shear strain stiffening analysis, once again, we performed large amplitude oscillatory shear tests under varied levels of axial strain, correlated the collected mechanical data to compositional data of the uncompressed sample, and scaled in a similar manner as before.

To first determine if strain stiffening differs under different amounts of compression, we attempted to scale and collapse mechanical data for samples spanning bulk compressions from 5% to 49%, over ~ 8 different depths, with each sample tested under multiple applied compressive strains. We plot the collapsed data for all samples, all compressions, and all depths in Fig. 3a. We find that the scaled data collapse onto a single, universal curve for all compression levels. The high compression measurements extend the high-stress fitting range the closest to *γ/γ*_*c*_ = 1, constraining the power law exponent around 0.5. This collapse further illustrates the robustness of the mechanical phase transition associated with shear strain stiffening.

**Fig. 3.**
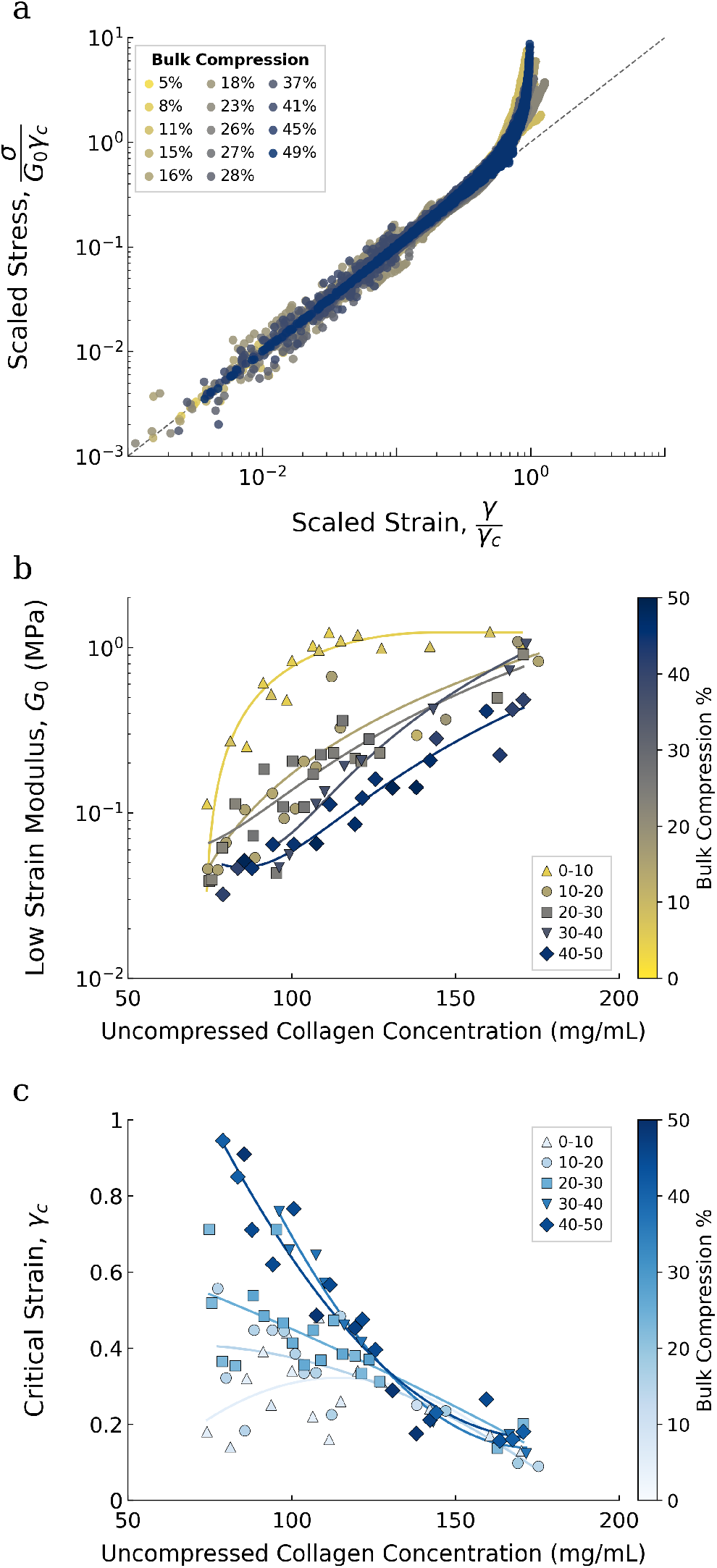
Analysis of mechanical and compositional properties under varying levels of static compression for N = 10 samples. a) Scaling collapse of depth-dependent mechanical data for ten different samples collected at bulk compressions ranging from 5% to 49% onto one universal curve. b) *G*_0_ and c) *γ*_*c*_ as a function of collagen concentration for different levels of compression. Local mechanical data were correlated with compositional data at the matching location in the uncompressed tissue. Mechanical data are binned into groups of compressive strain and averaged over collagen concentration.

To relate strain-stiffening behavior to tissue composition, we spatially matched each compressed measurement location to its corresponding position in the uncompressed sample, enabling direct comparison with the FTIR-I data. We then compared mechanical responses across samples at similar collagen concentrations under different levels of applied compression. Specifically, we plotted the low-strain elastic modulus and the critical strain as functions of the collagen concentration at the matched (uncompressed) location (Fig. 3b,c). To more easily identify trends, we group the data extracted from the Lissajous curves for each depth into concentration bins of 3 mg/mL and applied compression bins of 0% to 10%, 10% to 20%, 20% to 30%, 30% to 40%, and 40% to 50%. As a guide to the eye, we then fit a polynomial for each set of binned compressions. We find that increasing the applied compressive load decreases the zero-strain modulus (Fig. 3b). This trend is in line with previous studies that show that cartilage is more compliant under increased axial strain (9, 18). Additionally, we find that the critical strain, *γ*_*c*_, increases with applied compressive loads for similar collagen concentrations (Fig. 3c), indicating that compressed tissue will strain further before stiffening. At large enough collagen concentrations, however, this effect disappears because axial strains are localized to the articular surface where collagen concentration is relatively low (see supplementary materials for local axial strain measurement). Collectively, these results illustrate that strain stiffening in cartilage is sensitive to the amount of applied axial compression. The tissue becomes softer at low shear strains and reaches larger critical strains *γ*_*c*_ before stiffening.

### Effect of compression in simulations

To test whether the network models capture the observed sensitivity of strain stiffening to applied compression, as demonstrated by the experiments, we investigated shear strain stiffening in simulated networks that have been dilated and subsequently compressed. We hypothesized that, since collagen networks are swollen with water in cartilage, the effect of compression might be to squeeze out some of this water and allow some of the collagen fibers to relax. Therefore, we modified our simulations to follow a four-step procedure. First, double-networks were randomly generated, as in the previous section, with a fully-occupied weak subnetwork and a bond occupation probability *p*_strong_ = 0.4 in the strong subnetwork (Fig. 4a,i). These networks were subjected to a dilation strain *γ*_dilate_ = 0.3 to represent the initial prestress present in hydrated cartilage (Fig. 4a,ii). This dilation strain was applied incrementally, slowly increasing the strain on the network and relaxing the node positions at each step. Next, we applied a uniaxial strain *γ*_compress_ along the *z* direction (Fig. 4a,iii). Again, this was applied incrementally. The maximum compression strain *γ*_compress_ was varied from 0 to 0.3. Finally, a shear strain sweep was performed in order to probe the nonlinear shear mechanics of the compressed networks (Fig. 4a,iv). The shear strain was varied from *γ*_*xy*_ = 10^*−*3^ up to 2 in logarithmically spaced increments. Results were averaged over 50 randomly generated networks for each value of *γ*_compress_. The networks strain stiffen, as shown by the nonlinear behavior of the scaled stress-strain curves shown in Fig. 4b. Here, the shear strain is rescaled by its critical value *γ*_*c*_, which was determined using the same procedure described above (see discussion of Fig. 2), and the shear stress is rescaled by *G*_0_*γ*_*c*_. Additionally, as these networks are prestressed, it is necessary to subtract off a prestress *σ*_0_. For each value of *γ*_compress_, we determine *σ*_0_ by performing a linear fit, *σ* = *σ*_0_ + *G*_0_*γ* to low strains *γ <* 10^*−*2^. The fit values of *G*_0_ were used to measure the low-strain shear modulus.

**Fig. 4.**
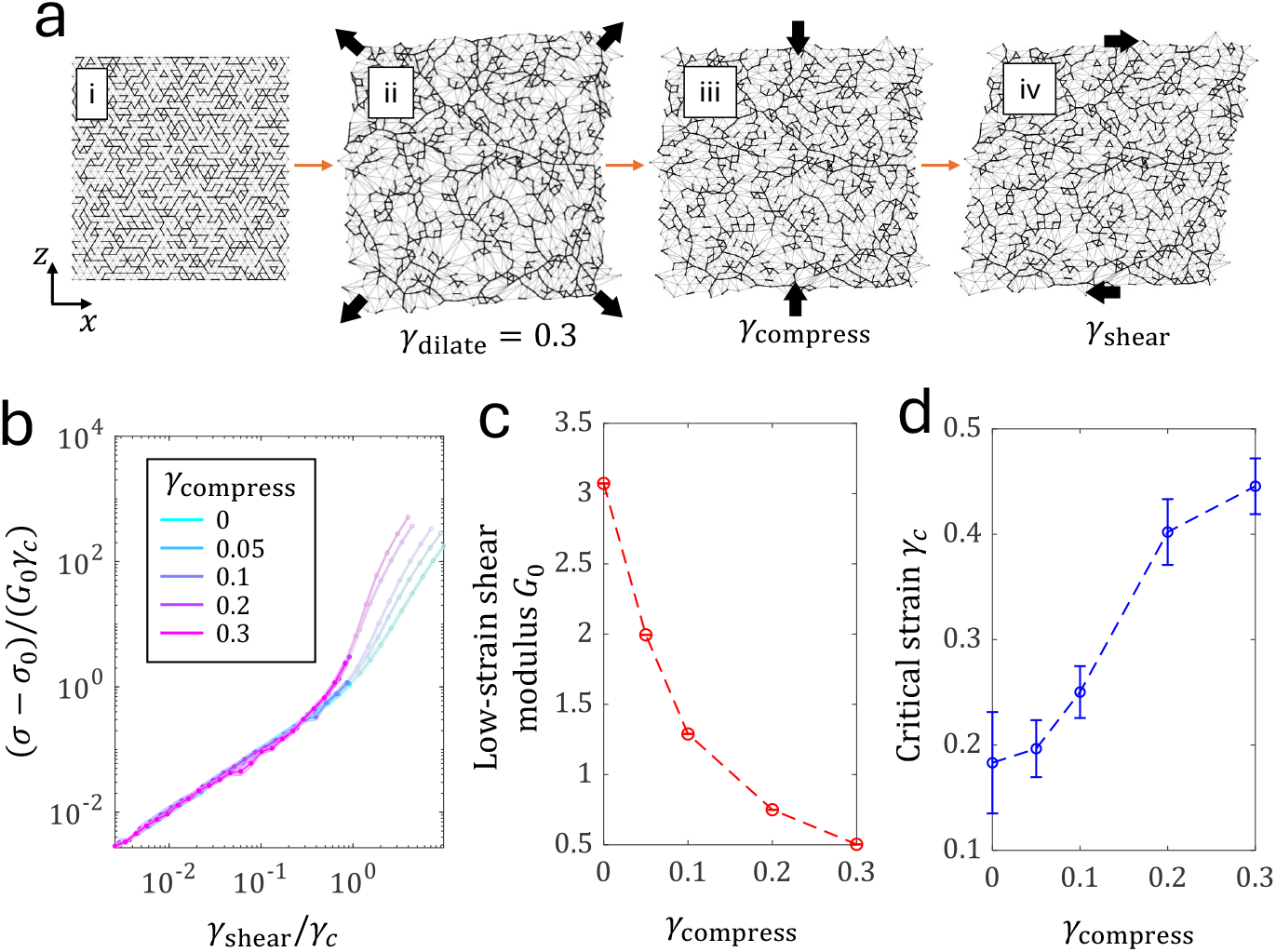
Network simulations reproduce the essential features of cartilage mechanics under multiaxial loading. (a) Schematic of simulation procedure in which (i) random networks are initially generated with a strong bond occupation probability of *p* = 0.4, then (ii) dilated, (iii) compressed, and (iv) subjected to a shear strain sweep. (b) Collapse of the shear stress-shear strain curves for various values of the compressive strain *γ*_compress_. (c) The low-strain shear elastic modulus as a function of *γ*_compress_. (d) The critical strain associated with strain stiffening as a function of *γ*_compress_.

We find that the curve collapse in the stiffening regime is not quite as robust as in the experiments. We hypothesize that the prestress protocol may have substantially stiffened the network prior to application of shear strain, which would change the expected slope of the transition region. Nevertheless, we find clear trends in the simulations that match the experimental results. Averaged across all simulation runs, *G*_0_ decreases with the compressive strain, as shown in Fig. 4c. This trend is consistent with what is observed in our experiments for low values of collagen concentration. In the network simulations, this compression softening can be understood mechanistically. When the networks are dilated, they undergo weak strain stiffening; the compression alleviates some of this pre-stress and therefore softens the networks. Finally, we observe that *γ*_*c*_ increases with *γ*_compress_, as shown in Fig. 4d, indicating that larger strains are needed for stiffening to occur in axially compressed networks. Thus, the network simulations reproduce the main effects observed in our experiments while providing simple explanations for how these effects could arise in cartilage.

## Discussion

Using experiments that measured how the local strain-stiffening response of cartilage depends on composition, and simulations of a network model suitably adapted for this study, we have established a framework to better understand the depth-dependent shear strain stiffening in this tissue. Prior work has established that this class of network models exhibits a mechanical phase transition from a regime where the network mechanics are governed by weak mechanical forces arising from effects such as bending of network elements to a regime where they are governed by stretching of network elements(47–49). To determine whether this framework applies to stiffening in cartilage, we ran parallel studies in the experiments and simulations. We find a number of corresponding results: first, the strain stiffening curves for different network compositions – collagen concentration in the experiments, and bond occupation probability in the simulations – collapse onto a universal curve; second, the linear elastic modulus increases for denser networks; third the critical strain at which the networks stiffen decreases for denser networks; finally, we find in both experiments and the model that under compression, the linear elastic modulus decreases while the critical shear strain increases.

These results provide evidence that can be used to determine which constitutive model best captures shear strain stiffening in articular cartilage. By experimentally establishing a connection between cartilage’s strain-stiffening behavior and its depth-dependent collagen concentration, we show that models that describe strain-stiffening as a result of the dynamics of an individual collagen fiber, such as the crimping model and the worm like chain (WLC) model, are insufficient in predicting the strain-stiffening response throughout cartilage tissue. In the case of the crimp model, the mechanics arise from a transition from straightening a crimped collagen fiber to the stretching of a straight collagen fiber. For the WLC model, the stiffening stems from the transition from random states to oriented states, and is dependent on thermal fluctuations. While fiber crimping and uncrimping may still play a role in shear mechanical behavior of cartilage, the WLC model is not applicable to collagen networks because the network is athermal, and so there should be no contributions from thermal fluctuations. In either case, because the response is dictated by the behavior of individual collagen fibers, changing the concentration of collagen would broaden the range but not change the mean critical strain of the tissue as a whole. Our measurements show that the critical strain does depend on the collagen concentration, and strongly suggest that these models do not capture the strain-stiffening behavior in cartilage tissue.

Furthermore, this study has shown that fiber realignment models of strain-stiffening also fail to capture key aspects of strain stiffening in articular cartilage. Fiber realignment models predict that the onset of stiffening is dependent on fiber orientation; highly aligned tissue should transition at smaller strains, while less aligned tissue requires higher strain to reorient the tissue and initiate the stiffening response. This mechanism predicts that the onset of stiffening depends on the distribution of fiber orientation and not the concentration of collagen fibers. The fact that only the upper ~ 50 µm of articular cartilage is characterized by highly aligned collagen fibers while the rest of the tissue is relatively isotropic strongly suggests that fiber orientation is not the main mechanism responsible for stiffening the bulk of the tissue. Fiber alignment models would suggest that this upper ~ 50 µm of the superficial zone should have a low critical strain. We note that because this region of the superficial zone is thin and very susceptible to compressive loads, the applied static compression drastically reduced the thickness of this highly aligned region to an indiscernible size. It would be informative to replicate this study under zero compression, to maintain the size of the aligned surface region, while simultaneously measuring fiber alignment to quantify and correlate collagen fiber orientation with the strain-stiffening response.

In contrast, we find that the 2-D network models *are* able to capture the dependency of strain stiffening on collagen concentration in articular cartilage. Network models describe strain stiffening as a transition from a “floppy” regime dominated by fiber bending forces to a “rigid” regime dominated by fiber stretching forces. The stiffening behavior depends on how close the collagen network is to rigidity. Changing the collagen concentration changes the distance of the network to rigidity, and so, network models predict a collagen concentration dependence where increasing the amount of collagen decreases the critical strain. This study confirms that prediction. Additionally, network models are able to capture the decrease of the linear shear modulus and the increase of the critical strain for stiffening under applied compressive axial strains. These results indicate that network models are able to capture the shear strain-stiffening behavior throughout the depth of the tissue. These correspondences provide strong evidence that stiffening in articular cartilage is primarily governed by a mechanical phase transition from a regime dominated by collagen fiber bending to one dominated by collagen fiber stretching.

We note that there are several limitations to our study. First, the mechanical data shown here are for neonatal bovine cartilage. While studies have shown the similarities of bovine tissue to human cartilage(19, 68), both age and species may play a role in tissue mechanics(78–81). Therefore, it would be instructive to replicate this study in human tissue to test whether the results found here hold in a different species and different age groups. The universality of the strain-stiffening behavior observed suggests that similar scaling analyses in other biological networks will also follow the same universal curve.

Second, although the network model used here is able to capture the behavior observed in the experiments, it is a simplified representation of a much more diverse and complicated system. Articular cartilage contains many more constitutive elements not considered here, such as the secondary aggrecan network, variations in the fibril diameters as a function of depth and chondrocyte maintenance and remodeling of the extracellular matrix. These properties may also play important roles in the strain-stiffening behavior of cartilage.

Third, while this work identifies the primary mechanism responsible for strain stiffening, it is likely that multiple mechanisms may be at play. For example, it is possible that crimping and fiber alignment models could be used in combination with 2-D networks to create multi-scale models. Such multi-scale models may more accurately mimic the extracellular matrix by considering more than one length scale as well as different dominant mechanisms at each length scale (82). Finally, incorporating these mechanisms into multi-phase models could be a useful tool for modeling the behavior of articular cartilage in multiscale simulations of joint mechanics.

Despite these limitations, this framework provides an intuitive understanding of why strain stiffening in cartilage is depth dependent. Near the superficial zone, where the collagen concentration is lower, the fiber networks are less dense so that larger strains are required to activate stretching modes. Conversely, in deeper regions where the collagen concentration is higher and the networks are more dense, such stretching modes become active at smaller critical shear strains. This framework also helps alleviate misconceptions that can arise in such studies. For example, compression of cartilage tissue leads to an increased concentration of collagen throughout the tissue, especially in the superficial zone. Naively, one might expect that increasing the concentration of collagen through compression should lead to an increase in tissue stiffness. Here, we find the opposite. Compression makes the tissue more compliant. Within the context of this network model framework, however, these results become easy to interpret. Compression does not alter the connectivity of the collagen network. Instead, the network topology remains the same, but the network elements are compressed into a smaller volume, which reduces the contributions from collagen fiber stretching to the tissue mechanics. Thus, when a compressed network is sheared, it is more compliant than in the uncompressed state, and a larger critical shear strain is needed to activate stretching modes in the fibers.

Such effects are particularly prominent in the superficial zone near the articular surface, which even in the uncompressed state is more compliant (lower *G*_0_ (18)) than the rest of the tissue, and can sustain larger shear strains prior to stiffening. Under an applied bulk compression the superficial zone shows the highest axial strain and the largest increase in the critical shear strain. These mechanisms for concentrating shear strain in the superficial zone under large axial and shear loading likely serve as protective mechanisms for cells deeper in the tissue(83, 84). Such insights may help inform how tissue damage arises and progresses in articular cartilage when this region is damaged and is no longer functional. Moreover, these insights should aid in the design and fabrication of biosynthetic replacements for damaged tissue(85, 86). More broadly, the simplicity of the network model and the universality of the observed scaling collapse suggest that this framework may help in understanding stiffening behaviors in other types of cartilage and other biological fiber networks across a diverse range of species.

## ACKNOWLEDGMENTS

We would like to thank the members of the Hospital of Special Surgery’s Orthopedic Soft Tissue Research group as well as the members of Center for Advanced Materials and Engineering in Orthopedics for helpful support and discussion about this project. This work was supported by the National Science Foundation grant NSF Directorate for Engineering, Contract: CMMI-2449148. This work was also supported by the NIH National Institute of Arthritis and Musculoskeletal and Skin Diseases, Contract: 5R21AR083064-02. This material is based upon work supported by the National Science Foundation Graduate Research Fellowship under Grant No. DGE-2139899. S.J.T. was supported by NSF DMR-2327094. Finally, this work made use of the Cornell Center for Materials Research Facilities supported by the National Science Foundation under Award Number DMR-1719875.

