## Supplementary Materials for "Shear Strain Stiffening in Cartilage is Governed by a Mechanical Phase Transition"

### Supplemental Materials

October 5, 2026

#### 1 Best fit to scaled experimental data

The scaling process suggests that this non-linear behavior can be predicted using only $G_0$ and $\gamma_c$, and gives us a relationship between stress and strain for the strain-stiffening transition:

$$\frac{\sigma}{G_0\gamma_c} = \Sigma(\frac{\gamma}{\gamma_c}) \tag{1}$$

where $\Sigma$ is a function of $\frac{\gamma}{\gamma_c}$. We can derive the function $\Sigma$ by assuming a form for the relationship between stress and strain. The chosen form contains a linear component to approximate the linear response of the tissue under small strain and a divergent component to approximate the strain-stiffening response at large strain.

$$\sigma = A\gamma - \frac{C}{\gamma_c^\delta} + \frac{C}{(\gamma_c - \gamma)^\delta} \tag{2}$$

Where $A = G_0 - \frac{C\delta}{\gamma_c^{\delta+1}}$, $C$ is a fitting parameter related to the magnitude of the contribution of the divergent term, and $\delta$ is the power law exponent that determines how the data diverges. If we use the form of $\sigma$ in Equation 2 and substitute it in Equation 1, we are able to determine the scaling function $\Sigma$:

$$\Sigma_r^\delta(x) = \frac{1}{1+r\delta}x + \frac{r}{1+r\delta}\left(\frac{1}{(1-x)^\delta} - 1\right) \tag{3}$$

where x is the scaled strain, $\gamma/\gamma_c$, and $r$ is defined as:

$$r \equiv C/A\gamma_c^{\delta+1} \tag{4}$$

In this function, $r$ is a universal quantity, insofar as the shape of the scaling function is universal. We find that the values of $C$ and $r$ vary with the chosen value of $\delta$. The function $\Sigma$ is fit using the scaled data from all samples and compression levels (seen in Fig. 3a of main text). The logarithms were used for weighting to account for the large range.

#### 2 Dependence of the low-strain modulus and critical strain on aggrecan

We investigated the influence of the secondary aggrecan network on the strain-stiffening behavior of articular cartilage by correlating the mechanical data to aggrecan concentration using the same process that was done for collagen concentration in Fig. 3b and c. As before, we grouped the data extracted from the Lissajous curves for each depth into concentration bins of 1 mg/ml and applied compression bins of 0% to 10%, 10% to 20%, 20% to 30%, 30% to 40%, and 40% to 50%. We fit a polynomial for each set of binned compressions as a guide to the eye. We plot the low-strain modulus $G_0$ and critical strain $\gamma_c$ versus aggrecan concentration in Fig. 1a and b. We find there is not a strong dependence of $G_0$ or $\gamma_c$ on aggrecan concentration for any compression range.

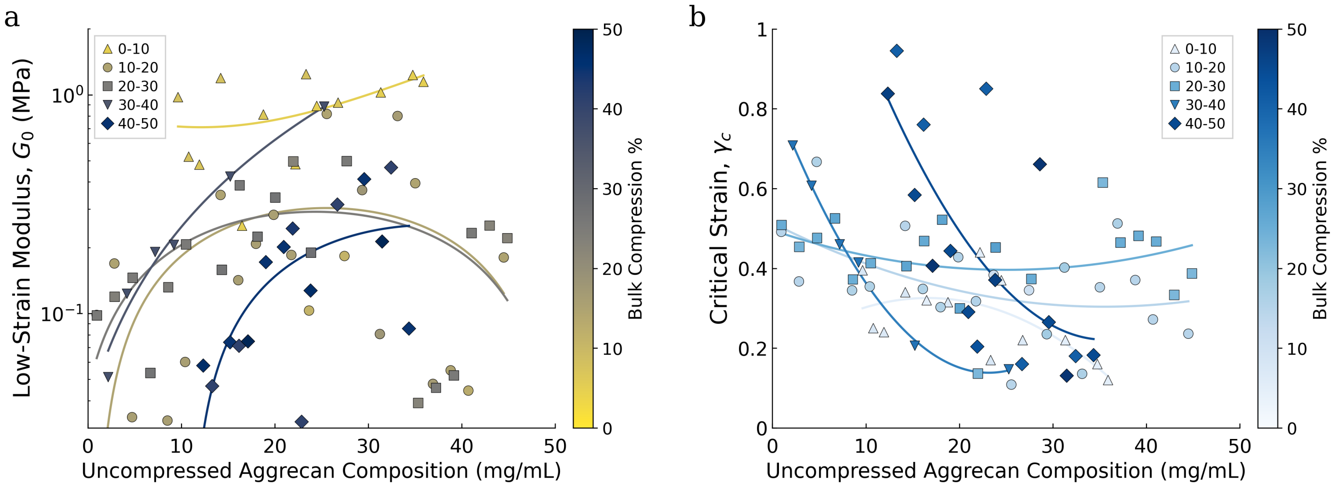


Figure 1: Analysis of mechanical properties under varying levels of static compression with respect to aggrecan concentration. a) $G_0$ and b) $\gamma_c$ as a function of aggrecan concentration for different levels of compression. Local mechanical data were correlated with compositional data at the matching location in the uncompressed tissue. Mechanical data are binned into groups of compressive strain and averaged over aggrecan concentration in the same manner done for collagen in Fig. 3b and c. of the main text

#### 3 Local Axial Strain Measurement

To correlate the mechanical data to the compositional data, we tracked prominent $\sim 8-12$ features of each sample before and after compression as seen in Fig. 2a. By finding the change in depth with compression for each feature, we determined the local axial strain for a given applied bulk compression level. We plot a representative sample in Fig. 2b. As expected, the tissue experiences greater axial strain closer to the articular surface. The amount of local strain flattens deeper in the tissue.

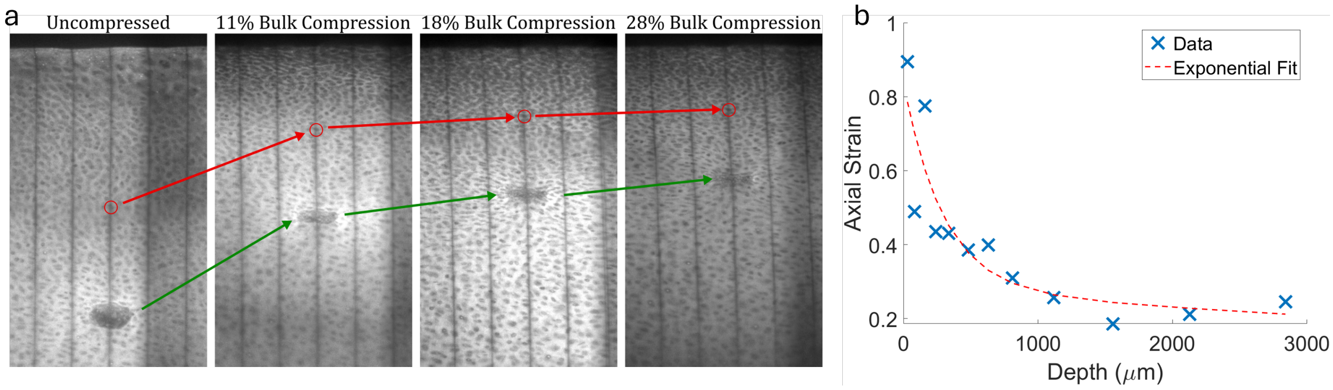


Figure 2: a) Tracking of two prominent features. The position of a chondrocyte is circled in red. The change in position of the cell is marked by the red arrows. The change of position of a blood vessel is marked by green arrows. b). For one sample, the local axial strain is plotted against depth in the uncompressed sample. The change in strain versus depth follows an exponential decay.

### 4 Scaling of the modulus in network simulations

For our sheared network simulations, we examined the scaling behavior of the shear modulus by plotting the differential shear modulus $G = d\sigma/d\gamma$ against the distance in strain to the critical strain $\gamma_c - \gamma$, as shown in Fig. 3. For each value of $p$, the modulus obeys a power law (*i.e.* a straight line on log-log axes) over some range of strains approaching the critical strain. Consistent with previous theoretical and numerical work (1, 2), the power law behavior does not persist for small values of $\gamma_c - \gamma$ (this would require the modulus to diverge at $\gamma_c$; in our model, the modulus remains finite). Furthermore, we observe a dependence of the scaling exponent on the value of $p$, ranging from roughly $-2$ at $p = 0.4$ to $-1$ at $p = 0.6$. The observation that this critical exponent depends on network topology (via the strong bond occupation probability) is consistent with the findings of other sheared network simulations, which show a range of non-mean-field exponents in 2D (3, 4) and in 3D (3, 5).

### 5 Effective medium theory for low-strain modulus

We perform an effective medium calculation for the small-strain behavior of a double network on a triangular lattice. As in the simulations described in the main text, we assume the lattice is occupied by a complete weak subnetwork with bond stiffness $k_w$ and a partially occupied strong subnetwork

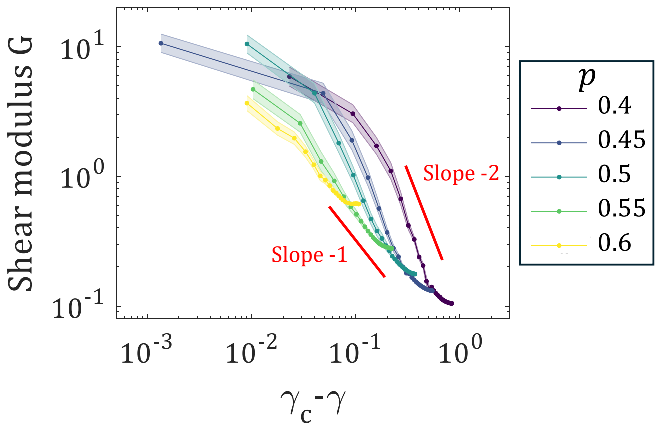


Figure 3: The shear modulus $G$ plotted against distance to the critical strain $\gamma_c - \gamma$ for the sheared network simulations. The modulus exhibits power law scaling with an exponent between $-2$ and $-1$, with a weak dependence on strong bond occupation probability $p$.

with bond occupation probability $p$ and bond stiffness $k_s$. In the effective medium theory, the strong subnetwork is replaced by a complete, softer network with bonds of stiffness $k_s^{\text{eff}}(p, k_w, k_s)$. This effective bond stiffness is chosen to approximate the linear response of the original diluted stiff subnetwork. The new complete subnetwork is combined with the weak subnetwork so that the effective stiffness of each bond in the joint lattice is predicted to be $k_w + k_s^{\text{eff}}(p, k_w, k_s)$. This enables a calculation of the combined double network's shear modulus.

#### 5.1 Triangular lattice calculations

We begin by computing the shear modulus per unit area of a complete triangular lattice with bond stiffness $k$. We consider a triangular lattice with one of its unit vectors pointing in the $x$ direction, subjected to a simple shear $\epsilon$ in the $x$ direction. We will compute the energy per unit area under this deformation, $E/A = C\epsilon^2/2$ where $C$ is the modulus per unit area. Each node has on average 2 associated diagonal bonds which stretch (horizontal bonds are not affected by the shear). Under shear, each diagonal bond changes its length from $a \to a(1 + \sqrt{3}\epsilon/4)$ to linear order in $\epsilon$. Therefore the energy per node is

$$\frac{E}{N^2} = 2 \cdot \frac{k}{2} \left( a \frac{\sqrt{3}}{4} \epsilon \right)^2 = \frac{3}{16} k a^2 \epsilon^2. \tag{5}$$

The area per node is a hexagon with an apothem of length $a/2$, which has an area of $a^2\sqrt{3}/2$. Therefore the energy per unit area is:

$$\frac{E}{A} = \frac{2}{a^2\sqrt{3}} \frac{E}{N^2} = \frac{\sqrt{3}}{8} k \epsilon^2. \tag{6}$$

Then, since $E/A = C\epsilon^2/2$, we identify the shear modulus per unit area $C = \left(\sqrt{3}/4\right) k$.

#### 5.2 Effective medium calculation

Next, we compute the effective stiffness of the complete network that approximates the linear response of the diluted stiff subnetwork. Call $D(q, k)$ the dynamical matrix of the triangular lattice with stiffness $k$ and wavevector $q$. The effective medium equation is

$$\frac{p - k_s^{\text{eff}}/k_s}{1 - k_s^{\text{eff}}/k_s} = \frac{1}{3}\frac{1}{s_{\text{BZ}}}\int_{\text{BZ}} \mathrm{d}^2 q \ \mathrm{Tr}\left[D\left(q, k_s^{\text{eff}}\right)\left(D\left(q, k_w\right) + D\left(q, k_s^{\text{eff}}\right)\right)^{-1}\right] \tag{7}$$

where the integral is performed over one Brillouin zone and $s_{\text{BZ}}$ denotes the area of a Brillouin zone (6–9). Now, note that $D(q, k) = k\, D(q, 1)$ because all bonds in the network have the same stiffness. This allows the integral to be done easily:

$$\frac{p - k_s^{\text{eff}}/k_s}{1 - k_s^{\text{eff}}/k_s} = \frac{2}{3}\frac{k_s^{\text{eff}}}{k_w + k_s^{\text{eff}}}, \tag{8}$$

where the factor of 2 comes from the trace of the $2 \times 2$ identity matrix. This equation may be written as a simple quadratic in $k_s^{\text{eff}}/k_s$ and solved, leading to:

$$\frac{k_s^{\text{eff}}}{k_s} = \frac{3}{2}p - \frac{3}{2}\frac{k_w}{k_s} - 1 + \sqrt{\left(\frac{3}{2}p - \frac{3}{2}\frac{k_w}{k_s} - 1\right)^2 + 3p\frac{k_w}{k_s}}. \tag{9}$$

The effective low-strain modulus per unit area for an $\epsilon$ simple shear is then:

$$C^{\text{eff}}\left(p, k_w, k_s\right) = \frac{\sqrt{3}}{4}\left(k_w + k_s^{\text{eff}}\right) = \frac{\sqrt{3}}{4}k_s\left(\frac{3}{2}p - \frac{1}{2}\frac{k_w}{k_s} - 1 + \sqrt{\left(\frac{3}{2}p - \frac{3}{2}\frac{k_w}{k_s} - 1\right)^2 + 3p\frac{k_w}{k_s}}\right). \tag{10}$$

Finally, for comparison with the results of simulations, the predicted shear modulus may be calculated by simply multiplying Equation 10 by the simulation area. The predictions of this effective medium theory are plotted along with the simulation data in Fig. 2f in the main text of this manuscript and seem to capture the trends in the simulations especially at the lower bond occupation probabilities $p < 0.55$. The disagreement at $p \geq 0.55$ arises from proximity to the strong subnetwork's rigidity percolation transition, which occurs at $p = 0.66$ (10). As the rigidity percolation transition is approached, the network's elasticity will include significant contributions from cooperative effects within the strong subnetwork, which are not accounted for by this effective medium theory.
